# Substrate stiffness regulates triple-negative breast cancer signaling through CXCR4 receptor dynamics

**DOI:** 10.1101/2025.03.28.645920

**Authors:** Kenneth K.Y. Ho, Johanna M. Buschhaus, Anne Zhang, Alyssa C. Cutter, Brock A. Humphries, Gary D. Luker

## Abstract

Biophysical properties of the extracellular matrix (ECM), such as mechanical stiffness, directly regulate behaviors of cancer cells linked to cancer initiation and progression. Cells sense and respond to ECM stiffness in the context of dynamic changes in biochemical inputs, such as growth factors and chemokines. While commonly studied as isolated inputs, mechanisms by which combined effects of mechanical stiffness and biochemical factors affect functions of cancer cells remain poorly defined. Using a combination of elastically supportive surface (ESS) culture dishes with defined stiffnesses and single-cell imaging, we report here that culturing cells on a stiff (28 kPa) versus soft (1.5 kPa) substrate increases CXCR4 and EGFR expression and promotes greater ligand-dependent internalization of CXCR4. In addition to increased CXCR4 expression, a stiff ECM also increases basal activation of Akt and ERK as well as signaling through these kinases in response to CXCL12-α and EGF and promotes migration of triple negative breast cancer (TNBC) cells. These data implicate receptor dynamics as a key mediator of Akt and ERK signaling as a mechanism for adverse effects of enhanced ECM stiffness on disease progression in TNBC.

## Introduction

Interactions with extracellular matrix (ECM) molecules regulate both normal and malignant cells. Cells sense and respond to the ECM through G protein-coupled receptors, tyrosine kinase receptors, integrins, and other molecules ^1,2^. Receptor-mediated interactions allow cancer cells to adapt to their immediate microenvironment, tuning intracellular signaling cascades that control survival, proliferation, and motility ^3,4^. Past studies show that the ECM stiffens as cancer progresses, and both ECM alignment ^5^ and mammographic density ^6^ predict poor prognosis in breast cancer. Furthermore, other environmental inputs, such as growth factors, regulate cell physiology critical for breast cancer initiation and progression. However, since research frequently focuses on independent effects of ECM mechanics or signaling, underlying mechanisms integrating ECM mechanics and biochemical inputs on breast tumor progression remain incompletely defined. Here, we combine changes in ECM mechanical stiffness and treatment with growth factors to identify how these properties coordinately regulate cell behavior.

To address how biochemical inputs regulate the behavior of tumor cells, we recently reported how conditioning cells with chemotherapeutics and/or cytokines potentiates signaling from CXCL12-CXCR4 and EGF-EGFR through PI3K/Akt/mTOR and Ras/Raf/MEK/ERK ^7,8^, two pathways central for breast cancer progression and metastasis. The ECM not only exposes cells to these same biochemical inputs by binding these ligands but also gives biophysical cues that significantly alter cellular behavior. For example, early studies demonstrated that exposure to a more rigid environment may promote cell adhesion, stemness, viability, and differentiation ^9,10^. Studies have since identified Akt and ERK signaling pathways as critical mechanotransducers underlying these behaviors, where activation of these signaling pathways promotes growth and stemness ^11,12^. Yet, the underlying mechanical-mediated mechanisms influencing Akt and ERK signaling dynamics in breast cancer remains poorly characterized.

Here, we investigated integrated effects of ECM mechanical stiffness and signaling through CXCR4 on activation of Akt and ERK in triple-negative breast cancer (TNBC). We focused on Akt and ERK signaling pathways because these kinases represent established integration points for environmental signaling inputs ^13,14^. We discovered that a stiff substrate increases both total and cell surface levels of CXCR4 and EGFR. Furthermore, we show that substrate stiffness shifts CXCR4 receptor dynamics, where a stiffer substrate enhances CXCR4 receptor internalization in response to CXCL12-α. Under baseline conditions, cells cultured on a stiff substrate had greater response to various growth factors (serum, EGF, and CXCL12-α), showing greater amplitude and duration of Akt and ERK activation than cells cultured on a soft substrate. Consistent with these data, we find that cells cultured on stiffer substrates migrated more than on softer substrates. Overall, our data demonstrate that ECM mechanics regulate cellular responses to biochemical signals mediated at least in part by changes in receptor localization and ligand-dependent internalization.

## Results

### Increased substrate stiffness increases CXCR4 and EGFR expression and surface localization

Breast cancers typically exhibit greater stiffness than normal breast tissues with intratumoral variations in mechanical properties regulating gene expression of cancer cells in tumor progression ^15^. The tumor microenvironment is rich in chemokines and cytokines, such as CXCL12 and EGF, that drive tumor progression and metastasis ^16–18^. While previous work has shown that EGFR is upregulated in cells cultured on a stiffer substrate ^19^, the combined effects of mechanical stiffness and biochemical stimuli on cell signaling remain incompletely understood, especially for CXCR4-mediated signaling. To address this gap in knowledge, we first investigated the effect of substrate stiffness on CXCR4 expression. We cultured SUM149 and patient-derived Vari068 cells on elastically-supported surface (ESS) dishes with defined stiffnesses (1.5 kPa and 28 kPa). ESS dishes represent a range of values that TNBC cells would typically encounter in a tumor environment ^20–24^ (**Figure 1A**). We coated ESS dishes with fibrinogen to provide cells a relevant ECM component found in breast tumors (**Supplemental Figure S1**). Using qRT-PCR and Western Blot, we found increased gene expression and total protein levels of CXCR4 in cells cultured on a stiffer substrate for both SUM149 and Vari-068 cells (**Figure 1B-C**). Aligning with previous reports ^25^, we also found increased expression of EGFR in cells cultured on a stiff substrate (**Supplemental Figure S2A**). To further support our findings, we performed gene set enrichment analysis (GSEA) of RNA sequencing data from two gene sets in the Gene Expression Omnibus (GEO) reporting TNBC cells cultured on soft versus stiff matrices (**Supplemental Table S1**) ^26,27^. In both of these datasets, we found upregulation of CXCR4 in each comparison of a stiff versus soft substrate (**Figure 1D**). As CXCR4 localizes to the plasma membrane, we used flow cytometry to quantify surface expression of CXCR4 in SUM149 and Vari068 cells. We found increased amounts of surface CXCR4 in SUM149 (**Figure 1E**) and Vari068 (**Figure 1F**) cells on stiffer versus softer substrate. Similarly, both SUM149 and Vari068 cells also displayed increased surface EGFR expression on a stiffer substrate (**Supplemental Figure S2B-C**). Together, these data demonstrate that higher substrate stiffness increases expression of both CXCR4 and EGFR.

**Figure 1.**
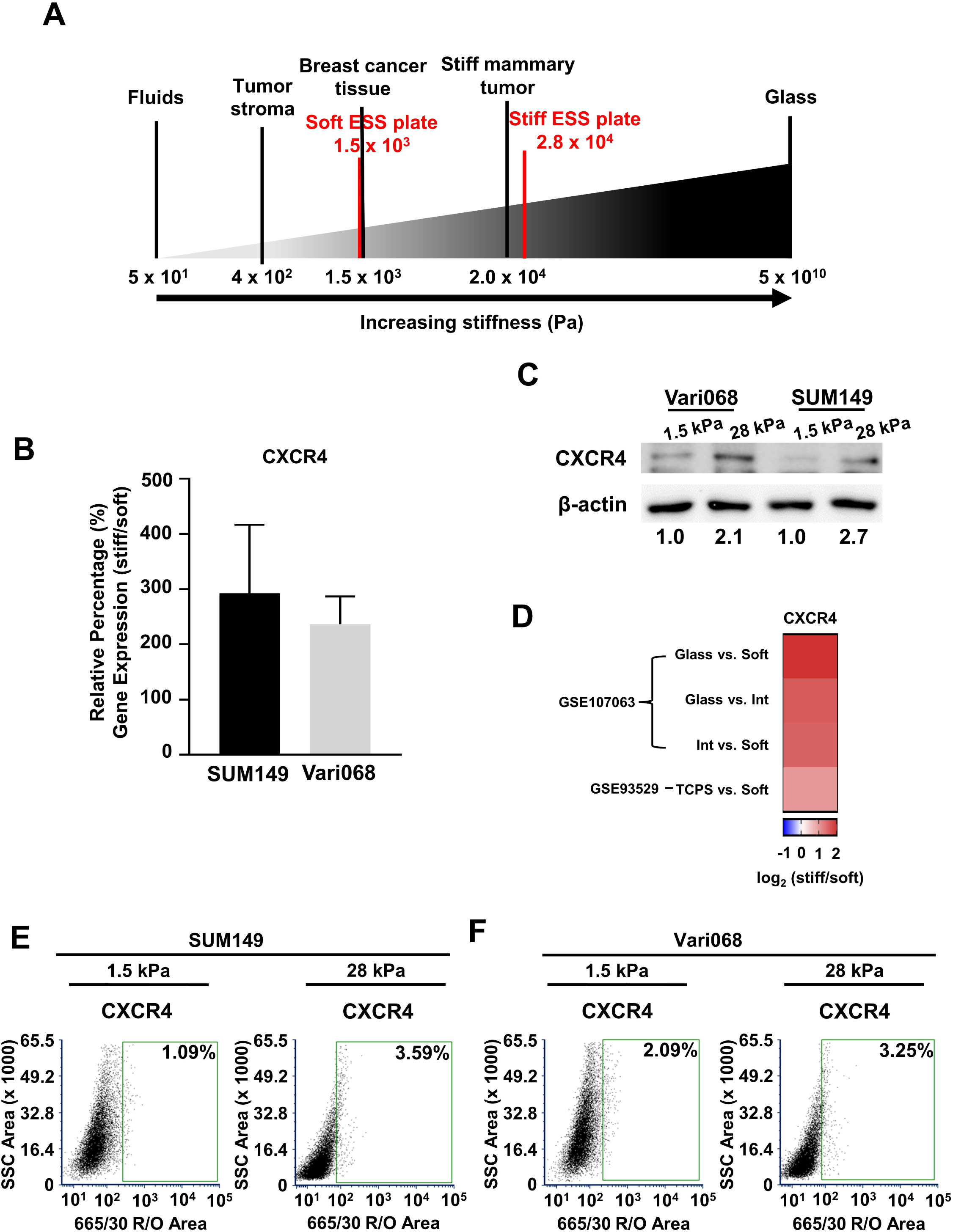
Enhanced substrate stiffness increases CXCR4 expression in triple-negative breast cancer. **A.** Schematic showing stiffnesses of the elastically supported surface (ESS) culture dishes compared to values found for primary breast tumors and the surrounding breast tumor environment ^20–24^. Fluids and glass are used as references. **B.** qRT-PCR analysis of CXCR4 in SUM149 and Vari068 cells grown on a stiff (28 kPa) substrate relative to cells grown on a soft (1.5 kPa) substrate, normalized to β-actin, showing higher CXCR4 expression in cells on a stiff substrate. Graphs show mean values ± SD (*n* = 6). **C.** Western blot analysis of CXCR4 levels in SUM149 and Vari068 cells cultured on a soft (1.5 kPa) or stiff (28 kPa) matrix. Values under the blots show values of CXCR4 normalized to β-actin for the same sample relative to cells cultured on a soft (1.5 kPa) matrix. **D.** Heatmap of CXCR4 with differential gene expression on stiff versus soft substrates in two genesets from Gene Expression Omnibus (GEO) accession numbers GSE93529 and GSE107063 ^26,27^. **E-F.** Flow cytometry plots for SUM149 (**E**) and Vari068 (**F**) cells show greater expression of CXCR4 on the cell surface in cells cultured on a stiff (28 kPa) versus soft (1.5 kPa) matrix (n ≥ 9,682 total cells analyzed in each condition). Cells stained with the IgG control antibody define the control gate.

Localization to the plasma membrane modulates availability of receptors for ligand binding and subsequent downstream signaling. To better visualize effects of substrate stiffness on CXCR4 localization, we imaged SUM149 and Vari068 cells expressing CXCR4-BFP (**Figure 2A**). Line intensity profiles showed higher fluorescence intensity from CXCR4-BFP at plasma membranes relative to the interior of cells (**Figure 2B**). Furthermore, we found increased percentages of fluorescence on cell membranes, defined as the percentage of quantifiable fluorescence at cell-cell junctions in an image, in cells cultured on a stiff substrate (**Figure 2C and Supplemental Figure S3**). We found similar results with EGFR, where a stiff substrate increased levels of receptor at the cell membrane (**Supplemental Figure S4A-B**) and higher percentage of fluorescence at cell membranes (**Supplemental Figure S3 and Supplemental Figure S4C**). Together these results demonstrate that increased substrate stiffness increases receptor localization of CXCR4 and EGFR at the plasma membrane.

**Figure 2.**
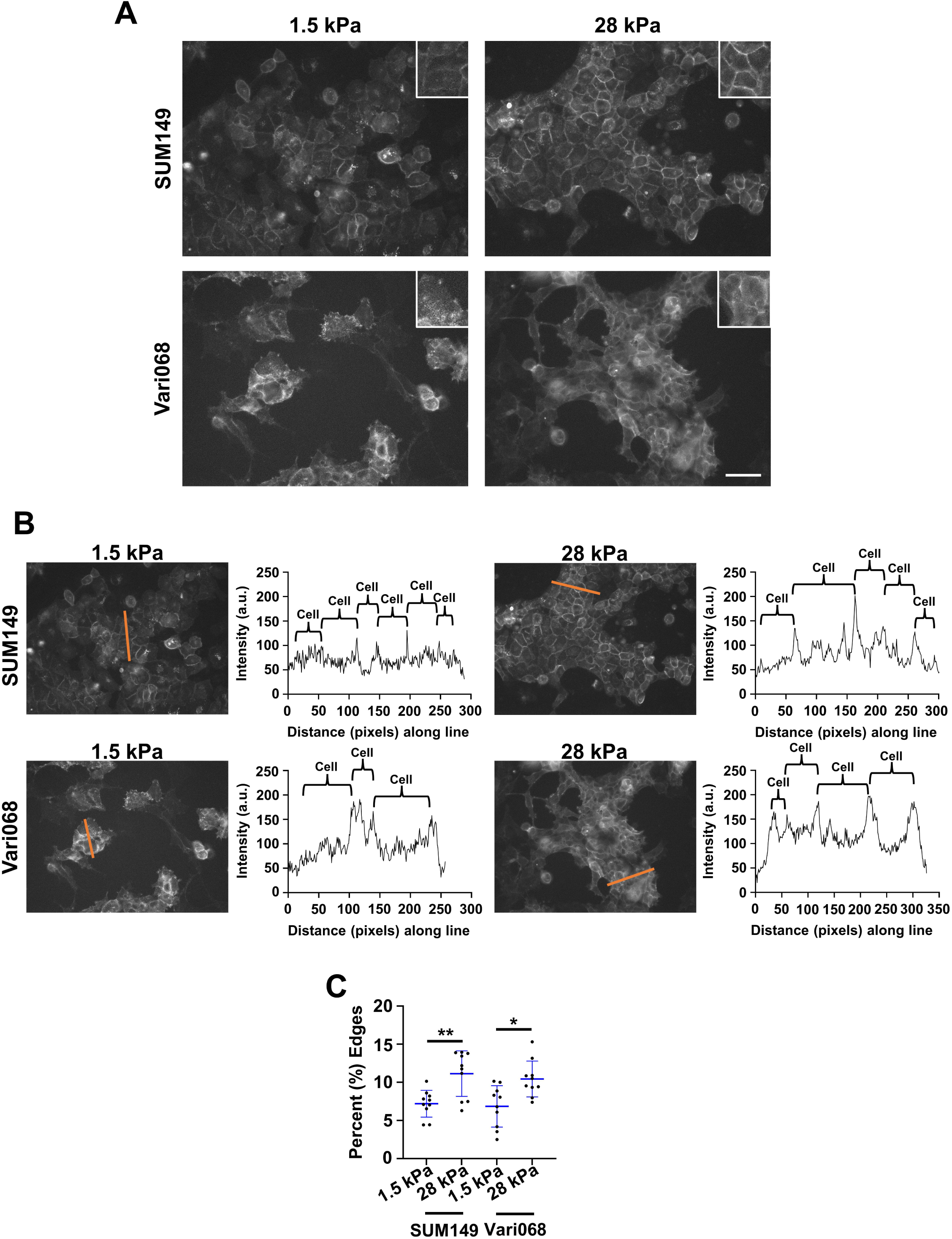
Enhanced matrix stiffness promotes altered CXCR4 localization. **A.** Representative images of SUM149 and Vari068 cells stably expressing CXCR4-BFP cultured on soft (1.5 kPa, **left**) and stiff (28 kPa, **right**) matrices under baseline signaling conditions described in methods. Scale bar is 50 µm. **B.** Representative line intensity and correlated histograms showing increased cell membrane intensity of CXCR4 in cells cultured on a stiff (28 kPa) matrix. **C.** Graphs show quantification of the cell membrane of CXCR4 in cells cultured on soft (1.5 kPa) versus stiff (28 kPa) matrices. *: p < 0.05, **: p < 0.01.

### A stiff substrate alters CXCR4 dynamics

The plasma membrane is not only a physical barrier between the cell and its environment but also a dynamic structure that responds and adapts to mechanical stimuli such as mechanical stiffness. Areas of high plasma membrane order facilitate receptor clustering and activation of signaling cascades ^28^. Therefore, we next analyzed effects of substrate stiffness on membrane order using laurdan, a validated fluorescent probe for membrane polarity. Both SUM149 and Vari068 cells cultured on a stiffer substrate had decreased membrane disorder and increased membrane order than the same cells on a softer substrate (**Figure 3A**), suggesting that a stiffer substrate promotes cell membrane organization that facilitates receptor signaling ^29,30^.

**Figure 3.**
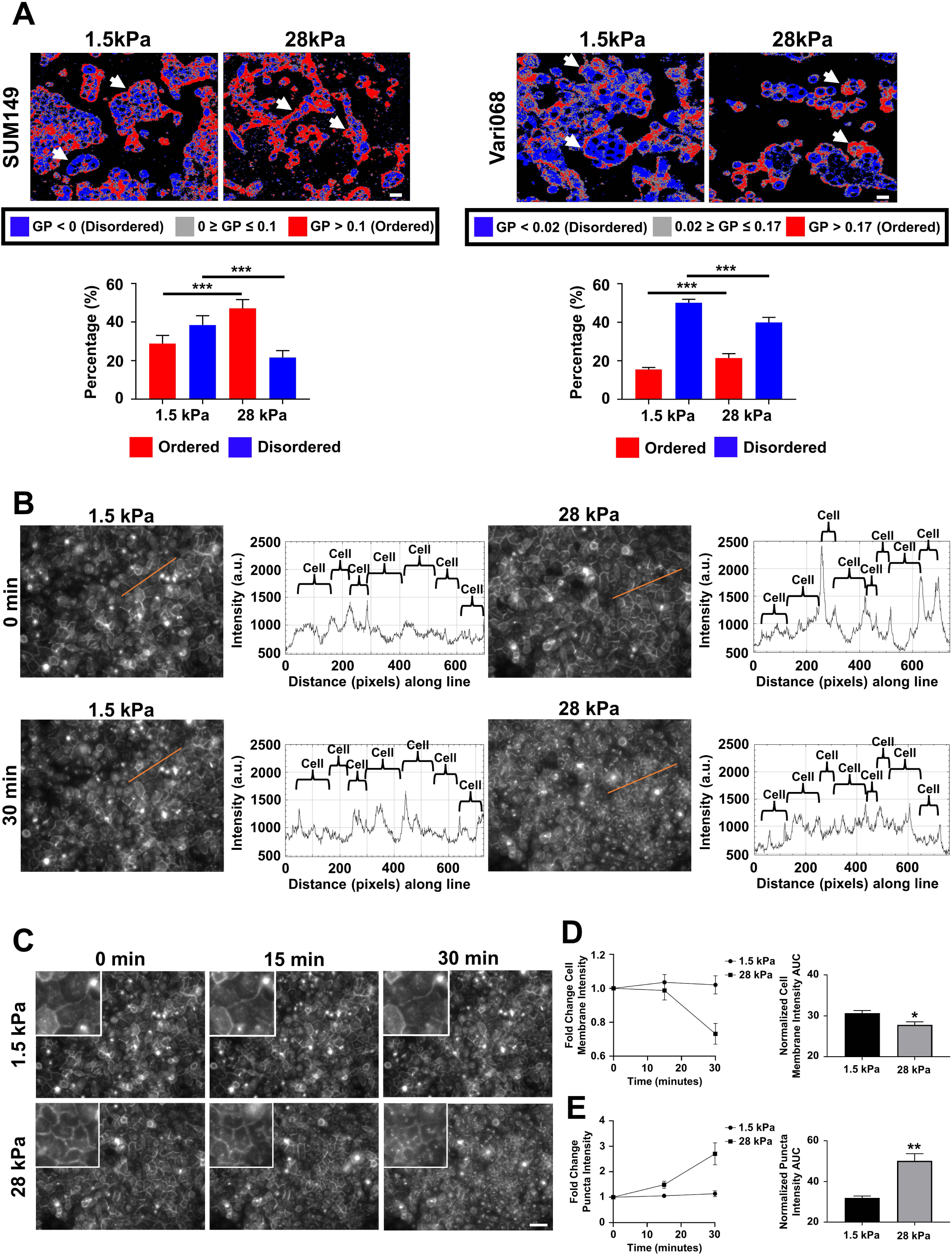
Greater ligand-dependent internalization of CXCR4 in cells cultured on a stiff matrix. **A.** Representative images of SUM149 or Vari068 cells stained with laurdan. Images show pseudocolored general polarization (GP) values. Red color indicates areas of high membrane order and less dynamics, whereas blue color denotes areas of low membrane order and more dynamics. Gray color indicates areas between the ordered and disordered phases based upon an arbitrary cutoff used for all groups. Arrows point to cell membranes. Scale bar□is□20□μm. Graphs show mean□+□SD for the percentage of all ordered and disordered pixels for each image above (n□=□10 images per group). ***: *p*□<□0.001. **B.** Representative line intensity and correlated histograms showing increased cell membrane intensity of CXCR4 in cells cultured on a stiff (28 kPa) matrix and a reduction in cell membrane intensity 30 minutes after addition of CXCL12-α (100 ng/mL) relative to that of cells cultured on a soft (1.5 kPa) matrix. **C.** Representative images of SUM149 cells stably expressing CXCR4-GFP cultured on soft (1.5 kPa, **top**) and stiff (28 kPa, **bottom**) matrices at initial time point (0 mins) and 15 and 30 minutes after addition of CXCL12-α (100 ng/mL). Images in **B** and from **C** come from the same experiment. Scale bar is 200 µm. **D-E.** Graphs show the fold change and area-under-the-curve for cell membrane intensity (**D**) and puncta from internalized CXCR4-GFP (**E**) in cells cultured on soft (1.5 kPa) versus stiff (28 kPa) matrices. Puncta in images represent normal endocytosis of CXCR4 resulting from endosomal accumulation. *: p < 0.05, **: p < 0.01.

In addition to receptor localization at the plasma membrane, receptor dynamics can drive downstream signaling and behavior. Therefore, we investigated changes in dynamics of CXCR4 in SUM149 and Vari068 cells cultured on substrates of different stiffness. We quantified internalization of CXCR4-GFP after adding CXCL12-α. Cells on a soft substrate did not have significant changes to the line intensity profile for fluorescence at the cell membrane comparing the initial time point versus 30 minutes after addition (**Figure 3B**). As seen previously (**Figure 2B**), cells on a stiff substrate had increased fluorescence intensity at the cell membrane compared with cells on a soft substrate (**Figure 3B**). Treatment with CXCL12-α caused internalization of CXCR4-GFP and loss of fluorescence at the cell membrane (**Figure 3B**). As a complementary approach, we next tracked CXCR4 fluorescence over time after treating cells with CXCL12-α. Cells on a soft substrate showed no change of cell membrane fluorescence intensity over 30 minutes after adding CXCL12-α (**Figure 3C and 3D**). Conversely, cells on a stiff substrate showed a significant decrease in cell membrane fluorescence intensity over time (**Figure 3C and 3D**). Consistent with this decrease, cells on a stiff substrate showed increased levels of internalized CXCR4-GFP as fluorescent puncta (**Figure 3C and 3E**), which represents normal endocytosis of CXCR4 after binding ligand. Together, these data suggest that increased substrate stiffness promotes localization of CXCR4 to the cell membrane with enhanced internalization after treatment with CXCL12-α, likely contributing to enhanced signaling in cells on a stiff substrate.

### Increased matrix stiffness enhances single cell Akt and ERK responses to exogenous stimuli

Enhanced receptor expression and altered dynamics may alter signaling cascades, leading to increased downstream kinase signaling activity. Therefore, we next asked how cells cultured on stiff versus soft substrates respond to external biochemical stimuli. To do this we used a kinase translocation reporter (KTR) system ^31^ in SUM149 and Vari068 cells to measure combined effects of mechanical and biochemical stimuli on kinase activities of Akt and ERK, two kinases downstream of CXCR4 and EGFR. KTRs utilize known downstream substrates specific for Akt or ERK. Phosphorylation of the substrate drives reversible translocation of the reporter into and out of the nucleus (**Figure 4A**). These cells also contain histone 2B fused to mCherry (H2B-mCherry) to facilitate automated image processing. To test coupled effects of mechanics and biochemical inputs, we cultured SUM149 cells on ESS dishes for five days; treated cells with a stimulus; and, using live-cell imaging, quantified KTR values for Akt and ERK (**Figure 4B**). SUM149 and Vari068 cells cultured on substrates with greater stiffness exhibited higher baseline activities of Akt (**Supplemental Figure S5**) and ERK (**Figure 4C-D**). Following stimulation with CXCL12-α, cells cultured on a stiffer substrate responded with greater activities of Akt and ERK than cells cultured on a softer substrate (**Figure 4E**). Consistent with these data, we found that cells on a stiffer substrate also signaled more in response to 10% serum (**Supplemental Figure S6**) and EGF (**Supplemental Figure S7**). Collectively, these data show that higher stiffness amplifies Akt and ERK signaling in response to biochemical stimuli like CXCL12 and EGF.

**Figure 4.**
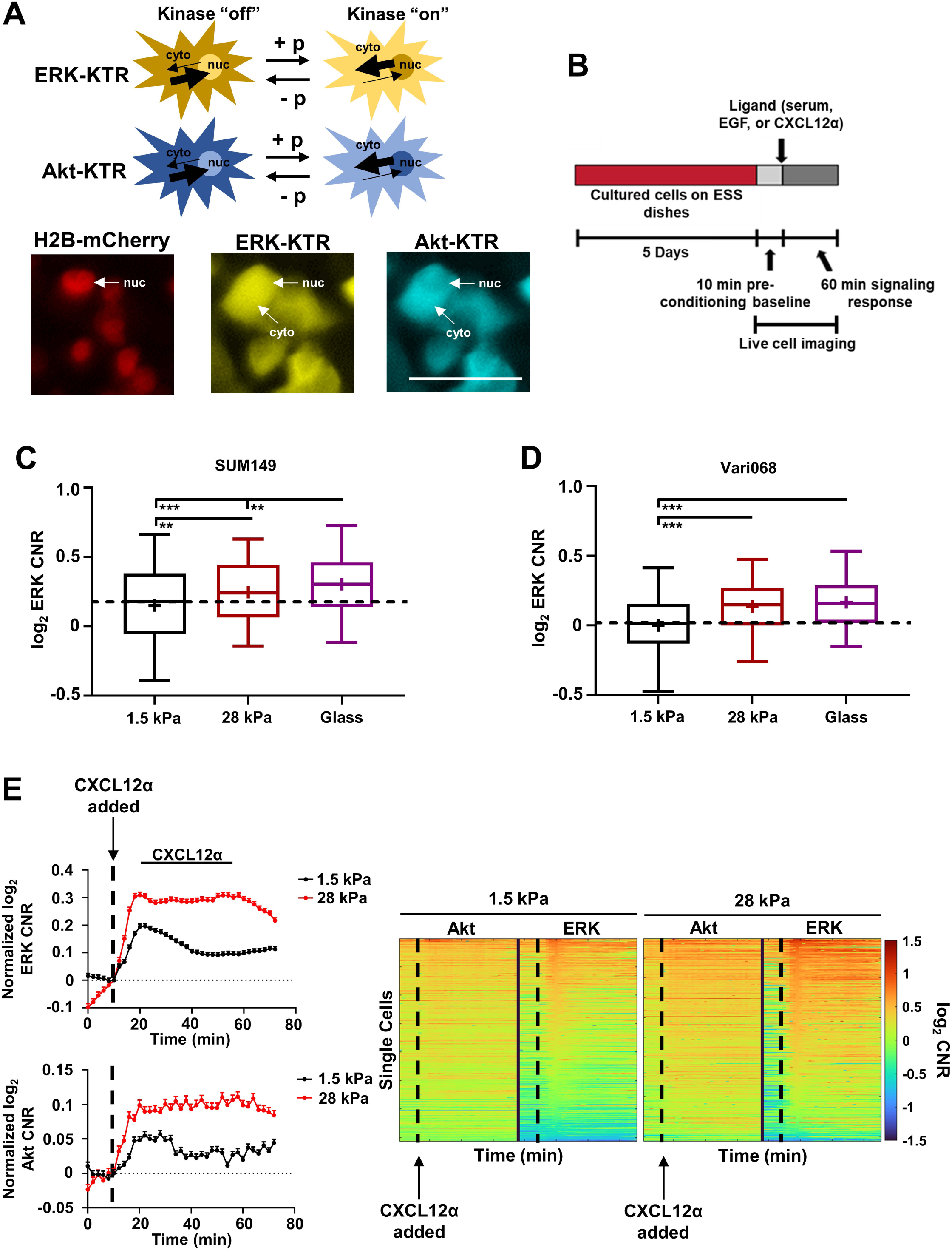
Enhanced matrix stiffness promotes single cell Akt and ERK signaling in response to stimuli. **A.** Schematic (**top**) and representative images (**bottom**) for action of ERK and Akt kinase translocation reporters (KTRs). When the KTR is phosphorylated (+ p), it accumulates in the cytoplasm (kinase “on”). Upon dephosphorylation (-p), the KTR translocates into the nucleus (kinase “off”). Translocation is reversible with continuous rather than binary readouts, allowing us to track dynamics of Akt (aquamarine) and ERK (mCitrine) in single cells. Scale bar is 50 μm. **B.** Representative schematic for single-cell imaging experiments. We cultured cells for 5 days on ESS dishes before acquiring single-cell, time-lapse imaging for 10 minutes before and 60 minutes after addition of stimulus. **C-D.** Box plot and whiskers for quantified log_2_ cytoplasmic/nuclear fluorescence intensities (cytoplasmic-to-nuclear ratio, CNR) for ERK (top) and Akt (bottom) activities in SUM149 (**C**) and Vari068 (**D**) at the time of making the wound in a monolayer of confluent cells (0 hr) (*n*□≥□180 cells per group). Line within the box denotes the median, and the “+” symbol denotes the mean. Dashed line represents the median of cells cultured on soft (1.5 kPa) ESS dishes. \**p* < 0.05, \*\**p*□<□0.01, \*\*\**p*□<□0.0001. **E.** We quantified activation of Akt and ERK in single SUM149 cells by imaging KTRs. Graphs show mean ± SEM for activation of Akt and ERK in an average cell in each condition in response to CXCL12-α (100 ng/mL) expressed as log_2_ of cytoplasmic to nuclear ratio (CNR) of fluorescence intensities normalized to the KTR value of the image before stimulus (t = 6) (*n*□≥□2,283 cells per group). Dashed vertical line denotes the time for adding a stimulus. **Right.** Single cell time tracks show activation of Akt and ERK in SUM149 cells quantified as the change in log_2_ CNR for each KTR in individual cells and displayed on a pseudocolor scale. A red color signifies increased Akt or ERK activity, while a blue color signifies decreased Akt or ERK activity. Compared to a soft environment, SUM149 cells cultured on a stiffer environment showed greater ERK and Akt activities in response to CXCL12-α. Dashed vertical line denotes the time point where the stimulus was added.

### Increased matrix stiffness enhances cell migration

Akt and ERK underlie cancer cell migration ^32,33^, and cancer cells experience a combination of external mechanical and biochemical stimuli during migration and metastasis. Therefore, we next investigated effects of mechanical stiffness on Akt and ERK signaling during migration on 1.5 kPa and 28 kPa substrates with glass as a supra-physiologic positive control. We discovered that greater matrix stiffness significantly increased migration of SUM149 cells (**Figure 5A and 5C**) with 28 kPa and glass producing the greatest effects. We observed similar results with Vari068 TNBC cells (**Figure 5B and 5D**). Additionally, cells showed comparable proliferation in each stiffness environment (**Supplemental Figure S8**), establishing that observed differences in wound closure mostly arose from effects on migration. We analyzed Akt and ERK activities at the time of wounding (0 hour) and the experimental endpoint (48 hours), stratifying cells into migratory (cells at the leading edge of the wound) or non-migratory (cells furthest from the leading edge in an image). At the time of wounding (0 hour), SUM149 (**Figure 5E, Supplemental Figure S9, and Supplemental Tables 3-4**) and Vari068 (**Figure 5F, Supplemental Figure S10, Supplemental Tables 5-6**) cells cultured on substrates with greater stiffness exhibited higher baseline activities of Akt and ERK. Furthermore, in all stiffness conditions ERK activity increased over time in both SUM149 (**Figure 5E, Supplemental Figure S9, Supplemental Tables 3-4**) and Vari068 (**Figure 5F, Supplemental Figure S10, Supplemental Tables 5-6**) cells. Both migratory and non-migratory cells at 48 hours showed greater activation of ERK than at the initial time point (0 hour), and migratory cells displayed higher ERK activity compared to the non-migratory population. Although SUM149 and Vari068 cells typically exhibit enhanced baseline activation of Akt due to loss/mutation of phosphatase and tensin homology deleted on chromosome 10 (PTEN) expression ^7,34^, we found that activity of Akt paralleled activity of ERK. Akt activity increased from 0 to 48 hours and migratory cells exhibited higher Akt activity than the non-migratory population in both SUM149 (**Figure 5E, Supplemental Figure S9, Supplemental Tables 3-4**) and Vari068 (**Figure 5F, Supplemental Figure S10, Supplemental Tables 5-6**) cells. Strikingly, Akt and ERK KTR values did not differ significantly among migratory cells on any of the stiffnesses tested, suggesting that mechanical stiffness primarily affects baseline Akt and ERK or signaling not related to migration.

**Figure 5.**
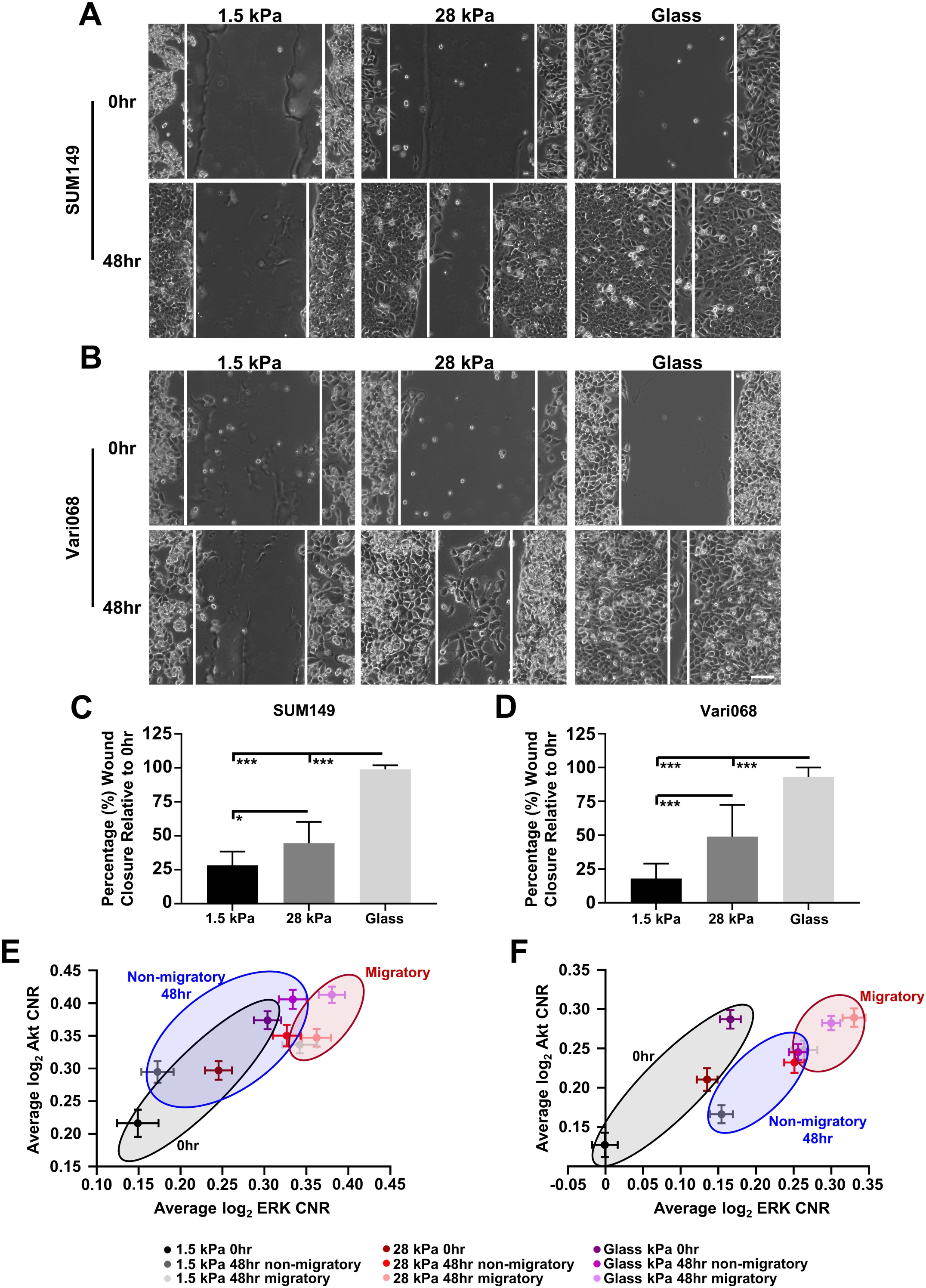
Enhanced substrate stiffness increases TNBC cell migration and kinase signaling. **A.** Representative wound healing images (n = 9) of SUM149 (**A**) and Vari068 (**B**) cells on ESS dishes with different stiffnesses at the time the wound was made (0 hr) and after 48 hr. Scale bar is 100 µm. **C-D.** Percentages of wound closure for SUM149 (**C**) and Vari068 (**D**) are presented as mean values ± SD (*n* = 12). *: p < 0.05, ***: p < 0.001. **E-F.** Graphs show mean ± SEM log_2_ CNR for Akt and ERK KTRs of SUM149 (**E**) and Vari068 (**F**) cells on different stiffnesses at time of the wound (0 hr) as well as migratory (at the leading edge) and non-migratory (furthest from the leading edge) cells at experimental endpoint (48 hr) (*n* ≥ 180 cells per group). Data show no significant difference between migratory cells in each group, suggesting changes to baseline Akt and ERK signaling regulate migratory capabilities on matrices with different stiffnesses.

## Discussion

Previous studies have defined a critical function of stiffness in tumor progression, where mechanotransductive signaling loops regulate almost all cellular behaviors ^35^. In this study, we provide evidence that increased mechanical stiffness facilitates promotes oncogenic behaviors of triple-negative breast cancer (TNBC) cells through CXCR4 receptor-mediated mechanism. Using elastically-supported surface (ESS) dishes, we found that enhanced substrate stiffness increased cell surface expression of CXCR4, aligned with increased expression of EGFR reported in past literature ^19^. Greater substrate stiffness increases membrane order and alters CXCR4 dynamics by promoting increased localization to the plasma membrane as well as increased internalization of CXCR4 to facilitate signaling. The combined effects of higher mechanical stiffness and biochemical stimuli produced enhanced and sustained Akt and ERK signaling at baseline and in response to specific ligands. With these changes, greater substrate stiffness also increases TNBC migration. These results highlight how ECM stiffness regulates dynamics and signaling of CXCR4 in TNBC cells.

During cancer progression, increased ECM deposition and crosslinking lead to a significant increase in tumor stiffness ^11^. Increased ECM stiffness can transmit physical signals to a cell, thereby changing behavior. Our data show that enhanced matrix stiffness increases membrane order. Areas of high membrane order, also known as lipid rafts, facilitate cell surface receptor activation, and disrupting these ordered membrane domains impairs downstream signaling cascades ^36,37^. An increase in ECM stiffness can cluster and increase expression of cell surface receptors, including integrins, receptor tyrosine kinases, and G protein coupled receptors ^11,38^. Consistent with our data, other studies also have shown increased CXCR4 and EGFR receptor expression on a stiffer substrate in other cell types ^25,39^. Therefore, we posit that increased membrane order leading to an increase in higher surface expression of CXCR4 and EGFR receptors. Furthermore, we speculate that cells sense ECM stiffness and respond by increasing CXCR4 and EGFR surface expression through i) coordination with integrins in these lipid rafts ^40,41^, and/or ii) the receptors themselves acting as mechanosensors ^39,42^, leading to a positive feedback loop. These potential mechanisms warrant further investigation.

Consistent with prior work ^43^, we showed that increased matrix stiffness enhanced cell migration. We also demonstrated that migratory cells exhibit higher activities of Akt and ERK than non-migratory cells. However, no significant differences existed for activities of Akt or ERK between migratory cells for any substrate stiffness. Since Akt and ERK signaling underlie cell migration, these results likely reflect that cells closer to the wound migrate more persistently than cells farther from the wound ^44^, which implies consistently elevated kinase signaling. We found the most significant differences in signaling at the initial time point, suggesting that the baseline effect of substrate stiffness on Akt and ERK signaling produces the major effects on migration.

Our results point to altered receptor internalization as a mechanism for how increased substrate stiffness drives CXCR4-mediated signaling. We found that increased substrate stiffness drives faster internalization of the CXCR4 receptor after binding ligand (**Figure 3C**), likely facilitating CXCL12α-CXCR4-mediated Akt and ERK activities through G protein and/or arrestin dependent signaling on endosomes ^45,46^. While it is unknown to what extent our results are specific to CXCR4 and our cell types versus a more general phenomenon across multiple cell types and receptors, other work has shown that increased substrate stiffness impacts ERK signaling through increasing the efficiency of EGFR activation and EGF membrane binding ^47,48^. Therefore, we suggest receptor localization and internalization as common mechanisms by which substrate stiffness drives oncogenic signaling.

Although all cells were exposed to the same conditions for each experiment, we note that coating ESS dishes with fibrinogen may impact the stiffness that cells experience. However, as previous work has shown that coating of PDMS with another matrix protein, collagen, had insignificant effects of stiffness ^49^, cells most likely experience the stiffness that the manufacturer provided.

In conclusion, this study demonstrates a critical link between ECM stiffness and signaling through two major receptors, CXCR4 and EGFR, in TNBC. Attempts to target enhanced tissue stiffness in other cancers have failed to provide patient benefits ^50,51^. More specifically, previous targeting of certain cell surface receptors such as integrins alone did not improve patient outcome ^52^. Instead, our data suggest that targeting, or co-targeting, changes in receptor localization and internalization benefit breast cancer patients by inhibiting key signaling processes needed for progression. Specifically, our data indicate that inverse agonists of CXCR4 that reduce steady state signaling by stabilizing the inactive form of receptor, such as T140 ^53^, or limiting CXCL12 binding could provide a method to limit TNBC progression. Overall, data from our study emphasize intersections of biochemical and mechanical inputs in determining localization and functions of both receptor tyrosine kinases and G protein coupled receptors in breast cancer.

## Materials and methods

### Cell Culture

We purchased SUM149 cells from the ATCC (Manassas, VA) and cultured cells in F-12 media supplemented with 10% fetal bovine serum (FBS), 1% penicillin/streptomycin (Pen/Strep) (Thermo Fisher Scientific, Waltham, MA), 1% GlutaMAX (Thermo Fisher Scientific), 5□μg/mL hydrocortisone, and 1□μg/mL insulin. We authenticated cells by analysis of short tandem repeats and characterized cells as free of *Mycoplasma* at the initial passage. Vari-068 cells (a gift from S. Merajver, University of Michigan) are patient-derived, TNBC cells adapted to cell culture. We cultured these cells in Dulbecco’s modified Eagle medium (DMEM) supplemented with 10% FBS, 1% Pen/Strep, and 1% GlutaMAX. We used all cells within 3□months after resuscitation and maintained all cells at 37□°C in a humidified incubator with 5% CO_2_.

### Vectors and cell lines

To identify activities of Akt and ERK in single cells, we used cells stably expressing fluorescent kinase translocation reporters (KTR) for these kinases as described previously ^7^. The KTR construct contains H2B fused to mCherry (H2B-mCherry) to denote the nucleus, an Akt-KTR (Aquamarine), ERK-KTR (mCitrine), and a puromycin selection marker all separated by P2A linkers and cloned into a piggyBac transposon vector (Systems Biosciences, Palo Alto, CA, USA). We transfected cells with the piggyBac transposon vector containing the KTR construct using FuGENE HD (Promega, Madison, WI, USA). For fluorescence imaging of CXCR4, we utilized a construct containing CXCR4 fused to a blue (CXCR4-mTagBFP2) or green (CXCR4-GFP) fluorescence protein ^7^. We selected stably expressing cells using puromycin (KTR) or flow cytometry (CXCR4-BFP and CXCR4-GFP) and confirmed expression by fluorescence microscopy ^54^. Only the breast cancer cells containing the KTR also stably expressed click beetle green luciferase (CBG; SUM149-CBG and Vari068-CBG) as described previously ^54^.

### Elastically supported surface (ESS) dishes

We utilized elastically supported surface (ESS) dishes (ibidi GmbH, Gr□felfing, Germany, 81199) to determine effects of a stiff (∼28 kPa), intermediate (∼15 kPa) or soft (∼1.5 kPa) environment on cellular behavior. These 35mm dishes have a 40 µm thick polydimethylsiloxane (PDMS) layer on top of a 100 µm thin glass cover slip to facilitate microscopy. To promote cell adhesion, we coated PDMS with fibrinogen per the manufacturer’s instructions. Briefly, we dissolved 100 µL of fibrinogen (Millipore Sigma, Burlington, MA, USA, 341576) or Alexa Fluor 488 conjugated fibrinogen (Invitrogen, F13191) (40 mg/mL stock concentration) in 4.9 mL of 1 M Tris-HCl ^55^ and added 800 µL directly to the PDMS. We incubated plates at 37°C for at least two hours; washed three times with phosphate-buffered saline (PBS); and then immediately plated cells onto the dish for cell experiments. For fluorescence imaging of fibrinogen coating, we used an Olympus IX73 microscope and DP80 CCD camera (Olympus) in the green channel using CellSens Software (Olympus).

### qRT-PCR

To analyze levels of the differentially expressed genes identified from transcriptomic data mining, we performed qRT-PCR using SYBR Green detection as described previously ^54^. Unless otherwise stated, we used and combined data from two unique sets of predesigned KiCqStart SYBR Green primers (Millipore Sigma) for the following genes:

CXCR4_1: 5’-AACTTCAGTTTGTTGGCTG-3’ and 5’-GTGTATATACTGATCCCCTCC-3’ CXCR4_2: 5’-CCTGAGTGCTCCAGTAG-3’ and 5’-AGATGATGGAGTAGATGGTG-3’

For the following gene, we used a single predesigned KiCqStart SYBR Green primer set (Millipore Sigma):

β-actin: 5′-TGTACGTTGCTATCCAGGCTGTGC-3′ and 5′-CGGTGAGGATCTTCATGAGGTAGTC-3′

### Flow cytometry and Western blotting

We seeded 8.0 x 10^4^ (Vari068) or 4.5 x 10^4^ (SUM149) cells in 800 µL Fluorobrite DMEM containing aprotinin as described above onto fibrinogen-coated PDMS in the center of an ESS dish. Two and four days after seeding, we changed dishes to fresh FluoroBrite DMEM containing 10% FBS. After overnight incubation (five days post seeding), we collected; stained; and analyzed cells by flow cytometry. In parallel dishes, we detected amounts of target proteins in total cell lysates by western blotting as described previously ^56^. We analyzed cell surface CXCR4 (Thermo Fisher Scientific, 17-9999-42) and EGFR (Cell Signaling Technology, Danvers, MA, USA, 5588S) using FCS Express for flow cytometry (De Novo Software, Pasadena, CA, USA). We used an IgG2a kappa-APC (Thermo Fisher Scientific, 17-4724-42) and IgG XP-Alexa Fluor 647 (Cell Signaling Technology, 2985S) for CXCR4 and EGFR isotype controls, respectively. For Western blotting we probed total cell lysates for CXCR4 (Santa Cruz Biotechnology, Dallas, TX, USA, sc-53534) or EGFR (Cell Signaling Technology, 4267), followed by β-actin (Cell Signaling Technology, 4967S) as a loading control. We quantified relative intensities of bands with ImageJ (National Institutes of Health, Bethesda, MD, USA).

### Transcriptomic data mining

For gene expression of cells cultured on stiff versus soft environments, we downloaded gene expression data from three different breast cancer studies ^26,27^ from the National Center for Biotechnology Information (NCBI) Gene Expression Omnibus (GEO) website. We took data from GSE107063, GSE83366, and GSE93529 datasets and identified common differentially expressed genes in at least two of the conditions. We analyzed RNA using iDEP, imported raw count matrices, and applied a filter using only genes with a minimum of one counts per million in at least half of the samples. We used rlog to transform counts data and treated missing values as zero. For individual genes, we defined significance as an adjusted *p*-value of less than 0.05 and absolute log_2_ fold change > 1. Finally, we performed DEseq2 with batch correction to quantify fold changes and p-values for each gene.

### Time-lapse single cell imaging, image processing, and fluorescence imaging of CXCR4

We quantified cellular localization of CXCR4 in cells that stably express CXCR4-BFP. We seeded 1.25 x 10^5^ SUM149 cells on fibrinogen-coated ESS dishes in FluoroBrite DMEM media containing 10% FBS, 1% Pen/Strep, 1% glutamine, 5□μg/mL hydrocortisone, 1□μg/mL insulin, and 0.01 U/mL aprotinin. Two and four days after seeding, we changed dishes to fresh FluoroBrite DMEM as described above containing either 10% (two days) or 1% (four days) FBS. After overnight incubation (five days post seeding), we imaged them by fluorescence microscopy as described above ^7^. We quantified localization of CXCR4 in cells using a line intensity profile in ImageJ. For CXCR4 dynamics, we seeded cells that stably express CXCR4-GFP into the center of an ESS dish and changed media on the dishes as described above. After overnight incubation (five days post seeding), we acquired multi-area, time lapse, z-stack images on an EVOS M7000 (Thermo Fisher Scientific). We captured images of only cells on the ESS at initial (0 min), as well as 15 minutes and 30 minutes after adding stimulus (final concentration, CXCL12 (100 ng/mL) or vehicle control). We used MATLAB to quantify CXCR4 localization at the cell membrane and as puncta in the cell. We first converted the z-stacked images into a 2D image using maximum intensity projection. Our program segments the intensity at the cell membrane and puncta using adaptive thresholding to detect the sharp increase in fluorescence intensity in the images. The program then separates cell membrane intensity and puncta by their distinct size and shape. The area and fluorescence intensity of the segmented cell membrane and puncta were calculated for comparison.

We quantified activation of both Akt and ERK kinases in single cells by fluorescence microscopy for KTRs. We acquired multi-area, time lapse images on an EVOS M7000 (Thermo Fisher Scientific). We captured images of only cells on the ESS every two minutes for six images before adding stimulus (final concentration, serum (10%), EGF (50 ng/mL R&D Systems), CXCL12 (100 ng/mL R&D Systems), or vehicle control) and then every two minutes for a total of 70 minutes. We calculated and displayed the ratio of median fluorescence intensities in the cytoplasm to the nucleus (CNR) as described previously ^7^.

### Laurdan staining

We analyzed effects of matrix stiffness on cellular membrane order with the fluorescent dye laurdan. We stained cells (5 µM laurdan, Cayman Chemical, 19706); imaged with an Olympus FV1000 RS multi-photon microscope; and calculated general fluorescence polarization (GP) of laurdan on cells cultured on a stiff or soft ESS dish as described in our previous publication ^54^.

### Wound healing assay

To determine effects of a rigid substrate on cell migration, we performed a wound healing assay. After coating ESS dishes as described above, we placed a 3-well insert (ibidi GmbH, 80369) in the middle of the coated dish and seeded 5.0 x 10^4^ SUM149 or Vari068 cells into each well before culturing overnight. The next day we removed the 3-well insert with forceps; washed with PBS; and added fresh FluoroBrite DMEM media (Thermo Fisher Scientific, A1896701), containing 10% FBS, 1% Pen/Strep, 1% GlutaMAX, and 0.01 U/mL aprotinin. For SUM149 cells, we supplemented medium with 5□μg/mL hydrocortisone and 1□μg/mL insulin. We acquired images immediately after making the wound (0 hour) and 48 hours later.

### Statistical analysis

For experiments comparing only two groups, we used a two-tailed, unpaired student’s *t* test. For experiments comparing multiple groups, we used one-way or two-way ANOVA where indicated and corrected for multiple comparisons with Tukey’s multiple comparisons test. We considered *p*□<□0.05 as statistically significant. We prepared bar graphs and dot plots (mean values + SD or SEM as denoted in figure legends) using GraphPad Prism 10.

## Acknowledgements

This study was supported by NIH grants R01CA238042, R01CA238023, R33CA225549, U24CA237683, R21AI173559, and R37CA222563. Research also was supported by funding from NSF award 2140104, DOD Breast Cancer Research Program W81XWH2210120, and the W.M. Keck Foundation.

## Author Contributions

K.K.Y.H. performed and analyzed experiments including single-cell imaging experiments; J.M.B. performed bioinformatics analysis; A.Z. performed and analyzed wound healing experiments; A.C.C. performed and analyzed flow cytometry experiments; B.A.H. and G.D.L. supervised and planned research; K.K.Y.H., B.A.H., and G.D.L. conceptualized study and wrote the manuscript; K.K.Y.H., J.M.B., A.Z., A.C.C., B.A.H., and G.D.L. revised the manuscript.

## Competing Interests statement

The authors declare no conflicts of interest.

## Data Availability Statement

The corresponding author (Brock A. Humphries) will provide data generated or methods used to analyze data upon reasonable request.

## Notes

### Competing Interest Statement

The authors have declared no competing interest.

