## Supplemental Figures for "Substrate stiffness regulates triple-negative breast cancer signaling through CXCR4 receptor dynamics"

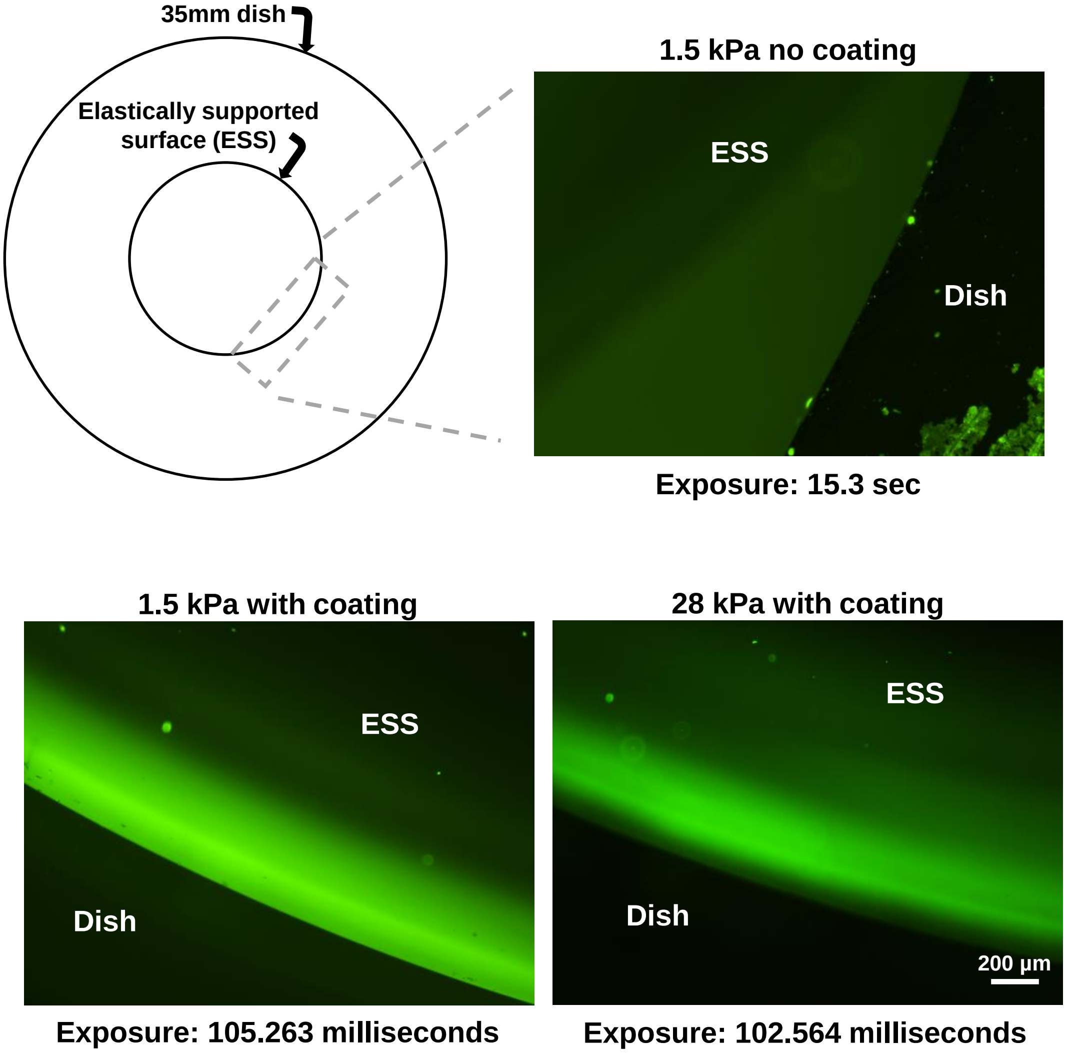


**Supplemental Figure S1. Coating of ESS dishes with fluorescent fibrinogen.** ESS dishes were coated with fibrinogen conjugated to AlexaFluor 488 as described in the Materials and Methods section to confirm successful coating and imaged after washing.

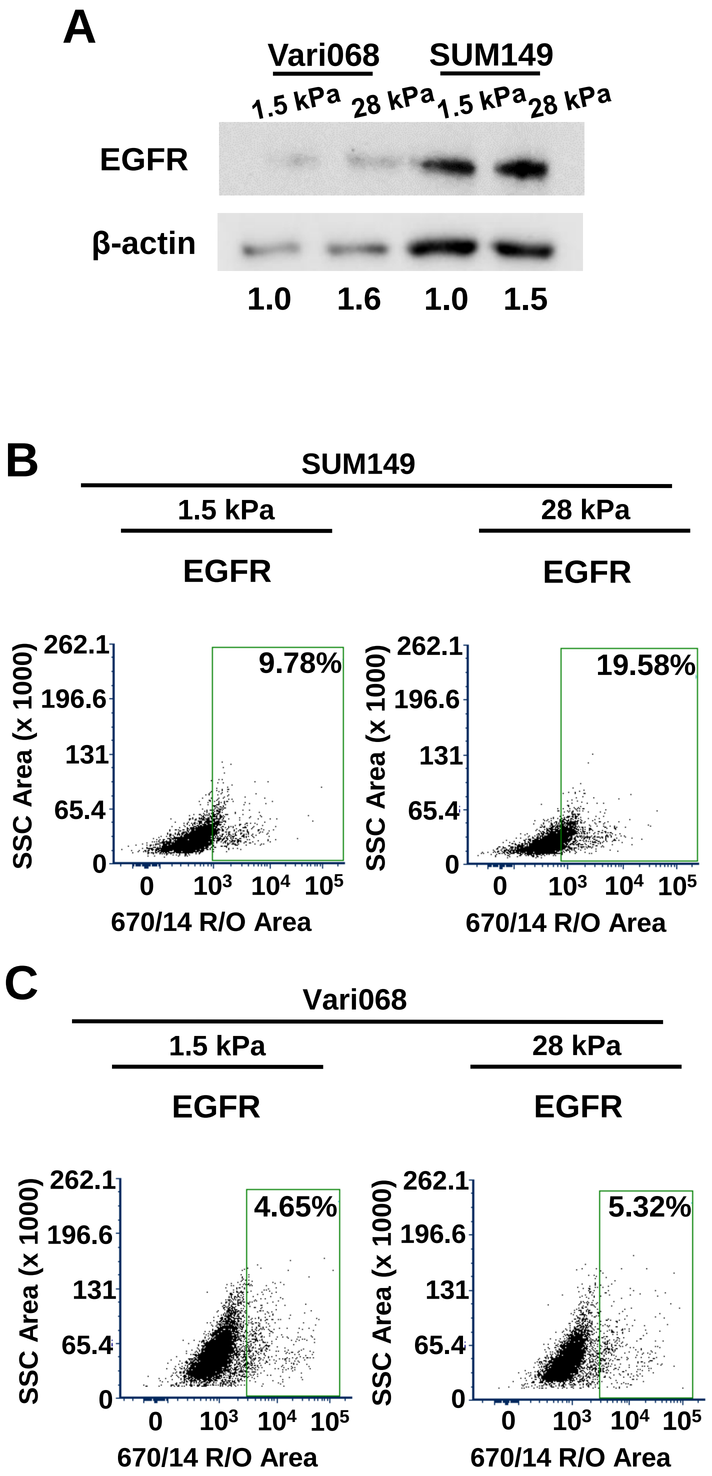


**Supplemental Figure S2. Increased expression of EGFR in cells cultured on a stiffer matrix. A.** Western blot analysis of EGFR levels in SUM149 and Vari068 cells cultured on a soft (1.5 kPa) or stiff (28 kPa) matrix. Values under the blots show values of EGFR normalized to β-actin for the same sample relative to cells cultured on a soft (1.5 kPa) matrix. **B-C.** Flow cytometry plots for SUM149 (B) and Vari068 (C) cells show greater expression of EGFR in cells cultured on a stiff (28 kPa) versus soft (1.5 kPa) environment. (n ≥ 9,433 total cells analyzed in each condition). Cells stained with the IgG control antibody define the control gate.

**Supplemental Table S1. Features of the three datasets used for bioinformatics analysis to determine drivers of enhanced Akt and ERK signaling on a stiff environment.**

| **Dataset** | GSE107063 | | | GSE93529 |
|---|---|---|---|---|
| **Platform** | Affymetrix array | | | Affymetrix array |
| **Cell Line Used** | MDA-MB-453 | | | MDA-MB-231 |
| **Conditions** | Fibronectin coated polyacrylamide gels; 5 and 30 kPa and glass | | | 0.7 kPa Hydrogels vs. TCPS |
| **Culture length** | 5 days | | | 2 days |
| **Replicates** | N = 4/condition | | | N = 4/condition |
| **Comparisons Used** | Glass v. Soft | Glass v. Mid | Mid v. Soft | Plastic v. Soft |
| **Genes UP** | 283 | 38 | 36 | 20 |
| **Genes DOWN** | 86 | 6 | 25 | 36 |

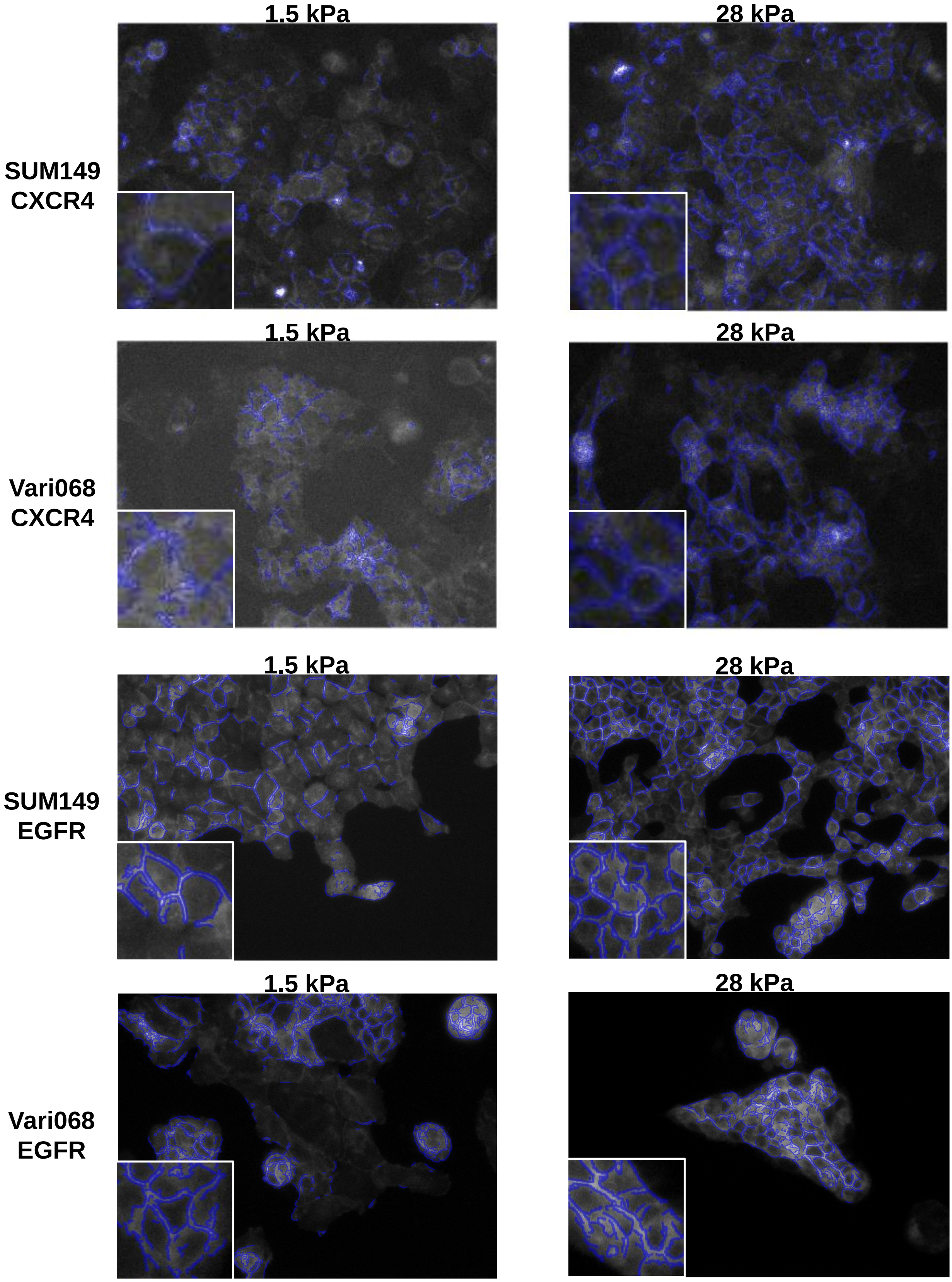


**Supplemental Figure S3. Cells cultured on a stiffer matrix have higher CXCR4 on cell edges.** Representative images of edge detection in SUM149 and Vari068 WT cells that stably express CXCR4-BFP for CXCR4 (**top**) and EGFR (**bottom**) cultured on 1.5 kPa (soft, **left**) and 28 kPa (stiff, **right**) matrices under baseline signaling conditions described in materials and methods. We analyzed cell edges with custom MATLAB code.

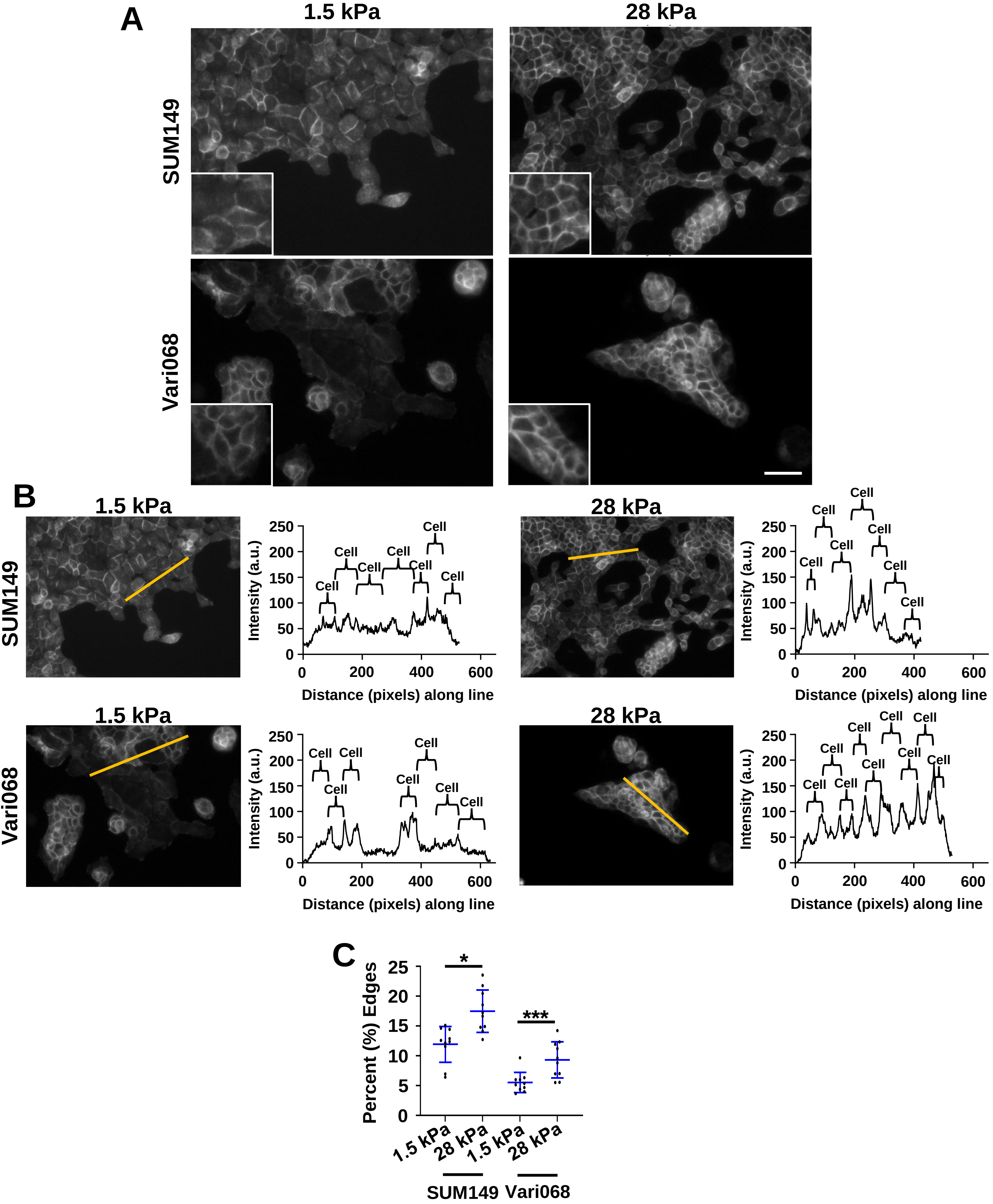


**Supplemental Figure S4. Cells cultured on a stiffer matrix have higher EGFR on cell edges. A.** Representative images of SUM149 and Vari068 WT cells stained with EGFR-AlexaFluor 647 cultured on 1.5 kPa (soft, **left**) and 28 kPa (stiff, **right**) matrices under baseline signaling conditions described in materials and methods. Scale bar is 50 µm. **B.** Representative line intensity and correlated histograms showing increased edge intensity of EGFR in cells cultured on a 28 kPa matrix. **C.** Graphs show quantification of the edges of EGFR in cells cultured on 1.5 kPa (soft) versus 28 kPa (stiff) matrices. *: $p < 0.05$, ***: $p < 0.001$.

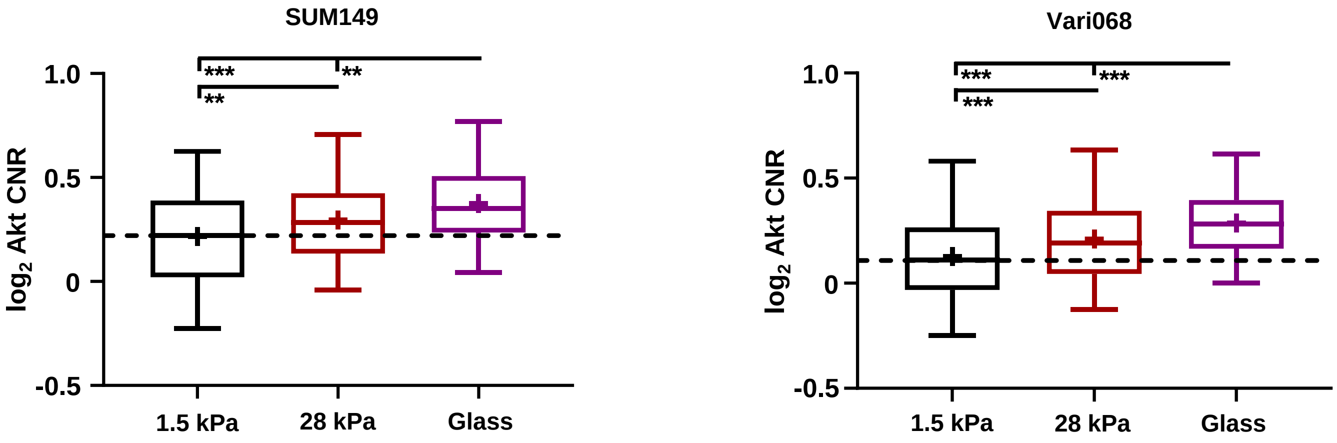


**Supplemental Figure S5. Enhanced substrate stiffness increased baseline Akt signaling in TNBC cells.** Box plot and whiskers for quantified $\log_2$ cytoplasmic/nuclear fluorescence intensities (cytoplasmic-to-nuclear ratio, CNR) for Akt activity in SUM149 (**left**) and Vari068 (**right**) at the time of making the wound in a monolayer of confluent cells (0 hr) ($n \geq 180$ cells per group). Line within the box denotes the median, and the "+" symbol denotes the mean. Dashed line represents the median of cells cultured on soft (1.5 kPa) ESS dishes. $^*p < 0.05$, $^{**}p < 0.01$, $^{***}p < 0.0001$.

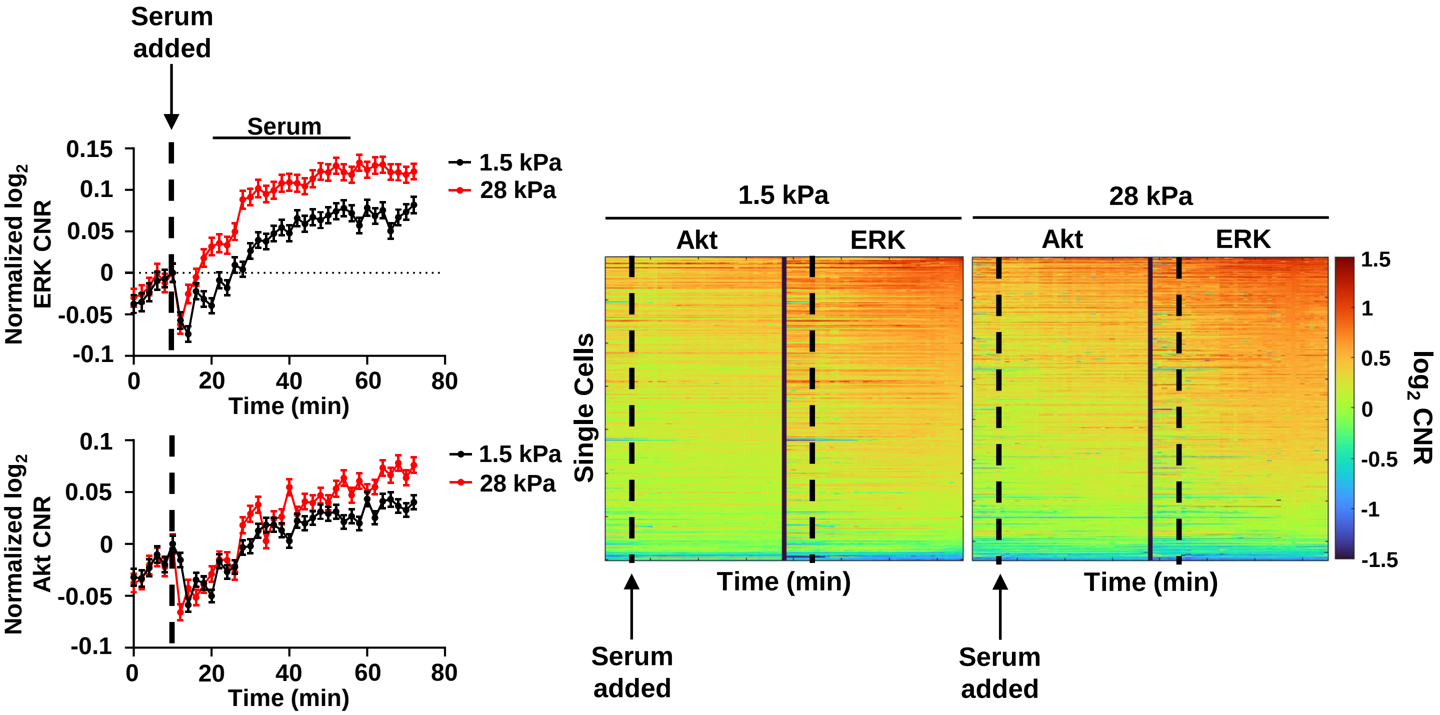


**Supplemental Figure S6. Enhanced matrix stiffness promotes single cell ERK signaling in response to serum.** We quantified activation of Akt and ERK in single SUM149 cells by imaging KTRs. Graphs show mean ± SEM for activation of Akt and ERK in an average cell in each condition in response to serum (10%) expressed as $\log_2$ of cytoplasmic to nuclear ratio (CNR) of fluorescence intensities normalized to the KTR value of the image before stimulus (t = 6) ($n \geq 2{,}283$ cells per group). Dashed vertical line denotes the time for adding a stimulus. **Right.** Single cell time tracks show activation of Akt and ERK in SUM149 cells quantified as the change in $\log_2$ CNR for each KTR in individual cells and displayed on a pseudocolor scale. A red color signifies increased Akt or ERK activity, while a blue color signifies decreased Akt or ERK activity. Compared to a soft environment, SUM149 cells cultured on a stiffer environment showed greater ERK signaling in response to serum. Dashed vertical line denotes the time point where the stimulus was added.

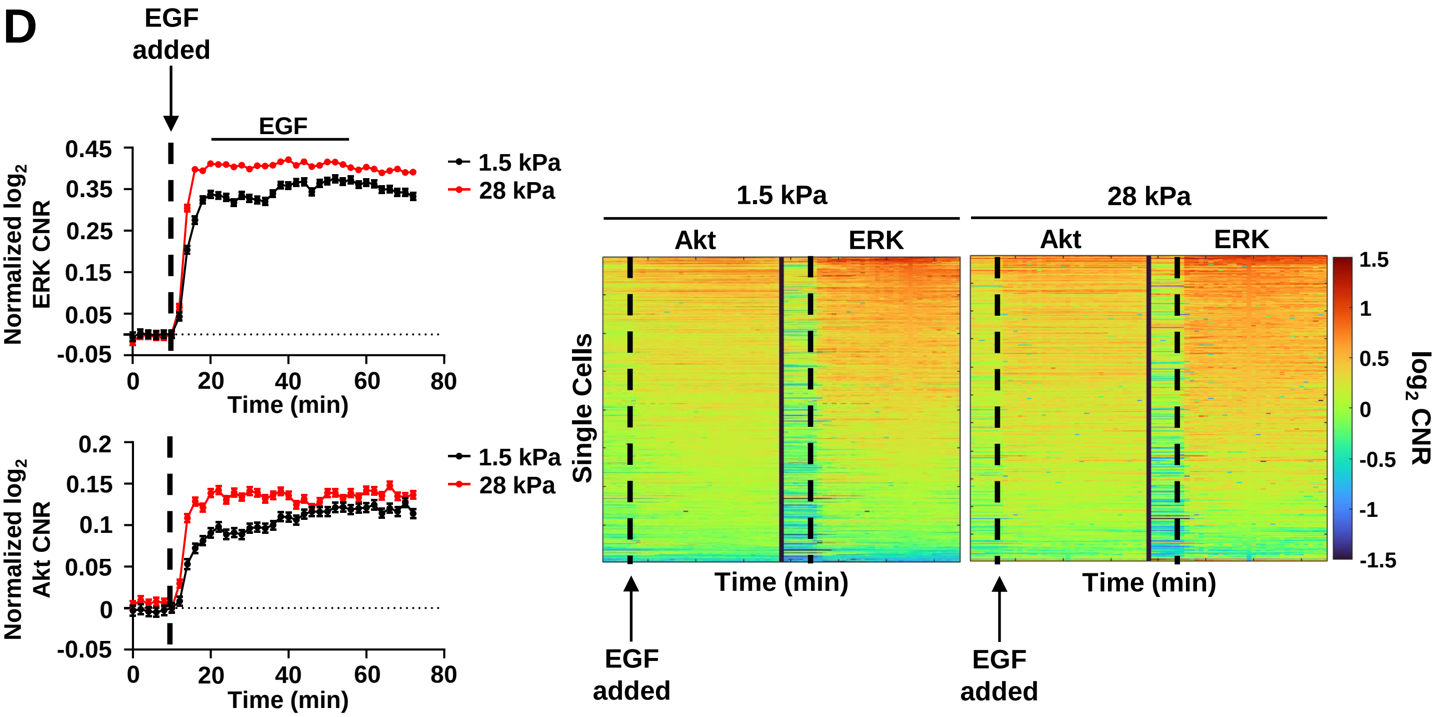


**Supplemental Figure S7. Enhanced matrix stiffness promotes single cell ERK signaling in response to EGF.** We quantified activation of Akt and ERK in single SUM149 cells by imaging KTRs. Graphs show mean ± SEM for activation of Akt and ERK in an average cell in each condition in response to EGF (50 ng/mL) expressed as $\log_2$ of cytoplasmic to nuclear ratio (CNR) of fluorescence intensities normalized to the KTR value of the image before stimulus (t = 6) ($n \geq 2{,}283$ cells per group). Dashed vertical line denotes the time for adding a stimulus. **Right.** Single cell time tracks show activation of Akt and ERK in SUM149 cells quantified as the change in $\log_2$ CNR for each KTR in individual cells and displayed on a pseudocolor scale. A red color signifies increased Akt or ERK activity, while a blue color signifies decreased Akt or ERK activity. Compared to a soft environment, SUM149 cells cultured on a stiffer environment showed greater ERK signaling in response to serum. Dashed vertical line denotes the time point where the stimulus was added.

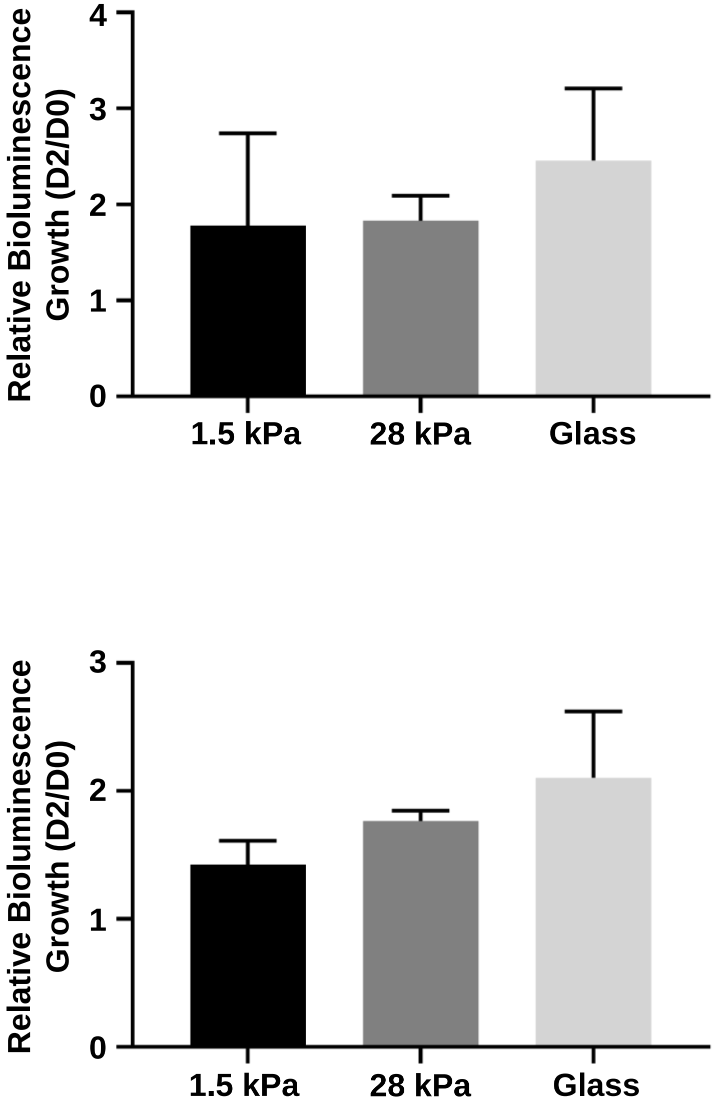


**Supplemental Figure S8. Substrate stiffness does not significantly change growth.** Relative growth of SUM149 (**top**) and Vari068 (**bottom**) cells on ESS dishes of 1.5 kPa or 28 kPa, or on glass. Graphs show mean values + SD for bioluminescence signal after 48 hrs (Day 2) relative to initial bioluminescence signal after seeding (Day 0) (n = 3).

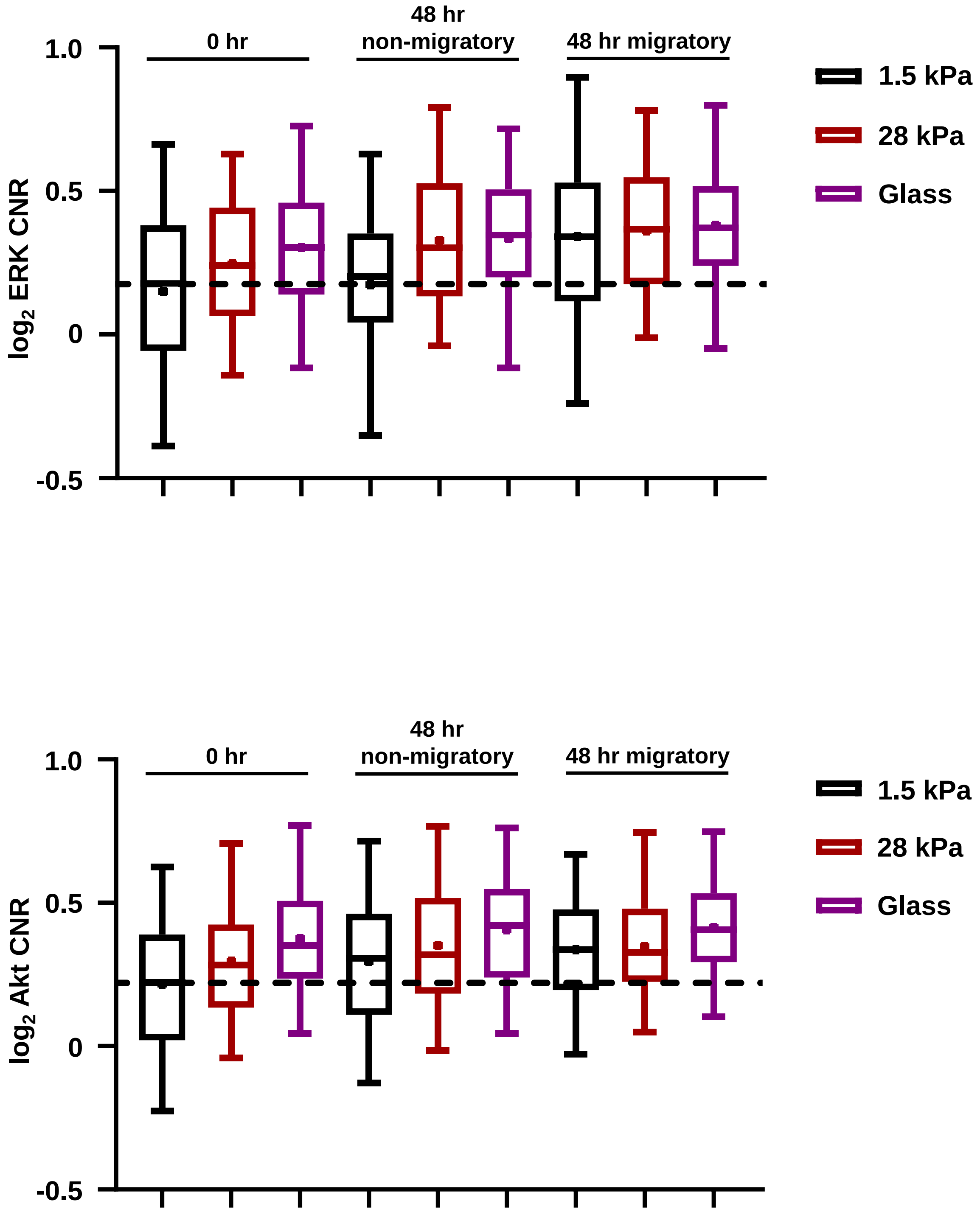


**Supplemental Figure S9. Increased matrix stiffness drives Akt and ERK signaling in SUM149 cells.** Box plot and whiskers for quantified $\log_2$ cytoplasmic/nuclear fluorescence intensities (cytoplasmic-to-nuclear ratio, CNR) for ERK (**top**) and Akt (**bottom**) activities based on the imaging from **Fig 2D**. Line within the box denotes the median, and the "+" symbol denotes the mean. Dashed line represents the median of the control group at the initial time point. See Supplemental Tables 1 and 2 for statistical comparisons.

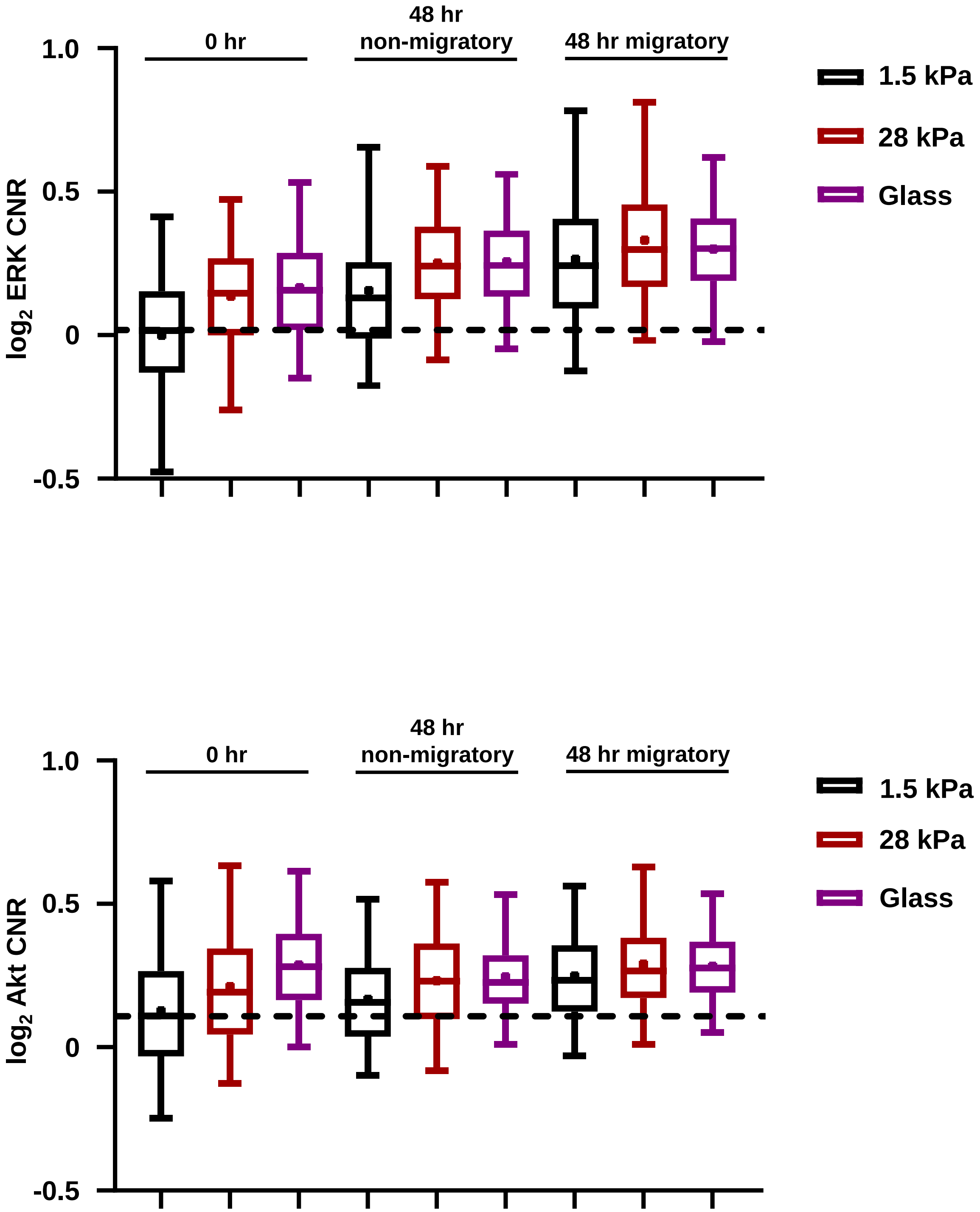


**Supplemental Figure S10. Increased matrix stiffness drives Akt and ERK signaling in Vari068 cells.** Box plot and whiskers for quantified $\log_2$ cytoplasmic/nuclear fluorescence intensities (cytoplasmic-to-nuclear ratio, CNR) for ERK (**top**) and Akt (**bottom**) activities based on the imaging from **Fig 2E**. Line within the box denotes the median, and the "+" symbol denotes the mean. Dashed line represents the median of the control group at the initial time point. See Supplemental Tables 3 and 4 for statistical comparisons.

**Supplemental Table S3. Calculated *p*-values for the ERK KTR in SUM149 cells from data in Supplemental Fig S9 using Tukey's multiple comparisons test.**

| | | 1.5 kPa | | | 28 kPa | | | Glass | | |
|---|---|---|---|---|---|---|---|---|---|---|
| | | 0 hr | 48 hr non-mig | 48 hr mig | 0 hr | 48 hr non-mig | 48 hr mig | 0 hr | 48 hr non-mig | 48 hr mig |
| 1.5 kPa | 0 hr | | | *** | * | *** | *** | *** | *** | *** |
| | 48 hr non-mig | | | *** | | *** | *** | *** | *** | *** |
| | 48 hr mig | | | | ** | | | | | |
| 28 kPa | 0 hr | | | | | * | *** | | * | *** |
| | 48 hr non-mig | | | | | | | | | |
| | 48 hr mig | | | | | | | | | |
| Glass | 0 hr | | | | | | | | | |
| | 48 hr non-mig | | | | | | | | | |
| | 48 hr mig | | | | | | | | | |

**Non-mig = non-migratory cell, mig = migratory cell, * = $p < 0.05$, ** = $p < 0.01$, *** = $p < 0.001$.**

**Supplemental Table S4. Calculated *p*-values for the Akt KTR in SUM149 cells from data in Supplemental Fig S9 using Tukey's multiple comparisons test.**

| | | 1.5 kPa | | | 28 kPa | | | Glass | | |
|---|---|---|---|---|---|---|---|---|---|---|
| | | 0 hr | 48 hr non-mig | 48 hr mig | 0 hr | 48 hr non-mig | 48 hr mig | 0 hr | 48 hr non-mig | 48 hr mig |
| 1.5 kPa | 0 hr | | * | *** | * | *** | *** | *** | *** | *** |
| | 48 hr non-mig | | | | | | | ** | *** | *** |
| | 48 hr mig | | | | | | | | * | * |
| 28 kPa | 0 hr | | | | | | | * | *** | *** |
| | 48 hr non-mig | | | | | | | | | |
| | 48 hr mig | | | | | | | | | |
| Glass | 0 hr | | | | | | | | | |
| | 48 hr non-mig | | | | | | | | | |
| | 48 hr mig | | | | | | | | | |

**Non-mig = non-migratory cell, mig = migratory cell, * = $p < 0.05$, ** = $p < 0.01$, *** = $p < 0.001$.**

**Supplemental Table S5. Calculated *p*-values for the ERK KTR in Vari068 cells from data in Supplemental Fig S10 using Tukey's multiple comparisons test.**

| | | 1.5 kPa | | | 28 kPa | | | Glass | | |
|---|---|---|---|---|---|---|---|---|---|---|
| | | 0 hr | 48 hr non-mig | 48 hr mig | 0 hr | 48 hr non-mig | 48 hr mig | 0 hr | 48 hr non-mig | 48 hr mig |
| 1.5 kPa | 0 hr | | *** | *** | *** | *** | *** | *** | *** | *** |
| | 48 hr non-mig | | | *** | . | *** | *** | | *** | *** |
| | 48 hr mig | | | | *** | | | *** | | |
| 28 kPa | 0 hr | | | | | *** | *** | | *** | *** |
| | 48 hr non-mig | | | | | | ** | ** | | |
| | 48 hr mig | | | | | | | *** | * | |
| Glass | 0 hr | | | | | | | | ** | *** |
| | 48 hr non-mig | | | | | | | | | |
| | 48 hr mig | | | | | | | | | |

Non-mig = non-migratory cell, mig = migratory cell, * = $p < 0.05$, ** = $p < 0.01$, *** = $p < 0.001$.

**Supplemental Table S6. Calculated *p*-values for the Akt KTR in Vari068 cells from data in Supplemental Fig S10 using Tukey's multiple comparisons test.**

| | | 1.5 kPa | | | 28 kPa | | | Glass | | |
|---|---|---|---|---|---|---|---|---|---|---|
| | | 0 hr | 48 hr non-mig | 48 hr mig | 0 hr | 48 hr non-mig | 48 hr mig | 0 hr | 48 hr non-mig | 48 hr mig |
| 1.5 kPa | 0 hr | | | *** | *** | *** | *** | *** | *** | *** |
| | 48 hr non-mig | | | *** | | * | *** | *** | *** | *** |
| | 48 hr mig | | | | | | | | | |
| 28 kPa | 0 hr | | | | | | *** | *** | | ** |
| | 48 hr non-mig | | | | | | | | | |
| | 48 hr mig | | | | | | | | | |
| Glass | 0 hr | | | | | | | | | |
| | 48 hr non-mig | | | | | | | | | |
| | 48 hr mig | | | | | | | | | |

Non-mig = non-migratory cell, mig = migratory cell, * = $p < 0.05$, ** = $p < 0.01$, *** = $p < 0.001$.

**Supplemental Table S1. Relevant GO Biological Processes Increased in Cells Cultured on Stiff Gels Based on GSEA.** We downloaded gene expression data from the National Center for Biotechnology Information (NCBI) Gene Expression Omnibus (GEO) website and analyzed data from GEO accession number GSE107063. Data show decreased signaling response and motility in cells cultured in soft compared to stiff environments.

| **GO Biological Process** | **p-value** | **q-value** |
|---|---|---|
| Response to hormone | 2.95E-20 | 2.15E-17 |
| Locomotion | 1.66E-17 | 6.03E-15 |
| Enzyme-linked receptor protein signaling pathway | 1.21E-15 | 4.04E-13 |
| Cell motility | 1.36E-15 | 4.09E-13 |
| Regulation of phosphorylation | 3.66E-15 | 1.01E-12 |
| Protein phosphorylation | 4.94E-15 | 1.32E-12 |
| Regulation of response to stimulus | 5.94E-15 | 1.55E-12 |
